# Single-cell analysis reveals a pathogenic cellular module associated with early allograft dysfunction after liver transplantation

**DOI:** 10.1101/2022.02.09.479667

**Authors:** Zheng Wang, Xin Shao, Kai Wang, Xiaoyan Lu, Li Zhuang, Xinyu Yang, Ping Zhang, Penghui Yang, Shusen Zheng, Xiao Xu, Xiaohui Fan

## Abstract

Liver transplantation (LT) is the standard therapy for patients with end-stage liver disease. Although LT technology has markedly progressed in recent decades, early allograft dysfunction (EAD) exacerbates the current organ shortage and impacts the prognosis of recipients. However, understanding of cellular characteristics and molecular events contributing to EAD is limited. Here, a large single-cell transcriptomic atlas of transplanted livers collected from four patients is constructed, including 58,243 cells, which are classified into 14 cell types and 29 corresponding subtypes with known markers, including liver parenchymal cells and non-parenchymal cells with different cell states. Compared to the pre-LT livers, graft remodeling is noted in the post-LT livers, with marked changes in several immune cells in either cell ratios or cell states. More importantly, an EAD-associated pathogenic cellular module is identified, consisting of mucosal-associated invariant T (MAIT) cells, granzyme B (GZMB)^+^ granzyme K (GZMK)^+^ natural killer (NK) cells, and S100A12^+^ neutrophils, all of which are elevated in EAD patient after LT. This cellular module is also verified in two independent datasets. Collectively, these results reveal the cellular characteristics of transplanted livers and the EAD-associated pathogenic cellular module at the single-cell level, offering new insights into the EAD occurrence after LT.

## INTRODUCTION

Liver transplantation (LT) is the most effective treatment for end-stage liver diseases, and the technology has made great progress over the past several decades.^1^ However, apart from the organ shortage as the major limitation of LT,^2^ early allograft dysfunction (EAD) following LT is considered a critical complication, with an incidence of approximately 20–40%, seriously affecting the long-term survival rate of allografts and recipients.^3–7^

EAD is a multifactorial complication of LT, and its risk factors include the donor risk index, surgery-related factors, and Model for End-stage Liver Disease score.^7–9^ Liver ischemia-reperfusion (IR) injury after surgery, a key factor for EAD development, is a complex process involving immune responses, inflammation, cell damage, and cell death, all of which are regulated by multiple cell lineages.^7, 10, 11^ However, the cellular heterogeneity, especially changes in the cell states or functions of immune cell populations, in complex liver diseases has restricted progress in understanding the detailed cellular characteristics and molecular events underlying the occurrence of EAD using conventional research methods.^12–14^

Recent advances in single-cell RNA sequencing (scRNA-seq) technologies have enabled the characterization of human liver tissues at high resolution,^12, 13^ with great advantages in the identification of novel cell types/states and molecular events involving complex physiological and pathological processes.^15–17^ In this study, we aimed to capture cellular heterogeneity using scRNA-seq and elucidate new cellular and molecular insights into EAD with this high-resolution approach. We constructed the largest single-cell transcriptomic atlas reported to date from four paired human transplanted livers before and after LT using scRNA-seq, and further performed module analysis to identify a pathogenic cellular module highly associated with EAD, which was further confirmed on two independent datasets. These will deepen our understanding of the cellular characteristics and pathogenic molecular events associated with EAD occurrence and provide a valuable reference for the prevention and treatment of EAD after LT.

## MATERIALS AND METHODS

### Human liver samples

All samples of transplanted livers were obtained from Shulan Hospital (Hangzhou, China) with Institutional Review Board approval (No. 2020065-77) and in conformance with the Helsinki Declaration (as revised in 2013). Written informed consent was obtained from the patients. No donor livers were obtained from executed prisoners or other institutionalized persons. Liver tissue samples were collected from four donors (before LT) with cold perfusion, and samples from the corresponding livers were collected from four recipient patients without hepatitis virus infection after 2 h of portal reperfusion.

### Collection of liver samples

After organ procurement, the allograft was fully perfused and preserved with 0–4°C Custodiol® HTK solution (Beijing, China). Before liver transplantation, 6 × 6 × 6 mm graft blocks were collected in tissue storage solution (10 mL, Miltenyi, Shanghai, China) as the preoperative samples (i.e., the ischemic specimen). Subsequently, the allograft was implanted using the piggyback technique, including the sequential procedures of inferior vena cava, portal vein, hepatic artery anastomosis, and biliary reconstruction. The postoperative samples (i.e., the reperfusion specimen) were collected just before closing the abdomen and stored in the same tissue storage solution (10 mL). All liver samples were subjected to the tissue dissociation process for preparation of single-cell suspensions.

### Preparation of liver single-cell suspension

The single-cell suspension was prepared with the human tumor cell isolation kit (Miltenyi) according to the manufacturer’s instructions. Briefly, the liver tissue was cut into pieces and placed into pre-prepared gentleMACS™ C tubes with dissociation enzyme mix. The appropriate gentleMACS™ program (37C_h_TDK_1) was then performed to dissociate the tissues for 1 h using the gentleMACS™ Octo dissociator with heaters. Subsequently, the cell suspension was filtered by a 40 μm nylon cell strainer (Corning, Shanghai, China) and transferred to a 50 mL centrifuge tube for centrifugation at 500 g for 3 min. The erythrocytes were lysed using 5 mL ACK lysing buffer (Gibco, Shanghai, China). Dead cells were removed with a Dead Cell Removal Kit (Miltenyi) according to the manufacturer’s recommendations. Finally, the cell pellet was washed twice and resuspended in 1 × PBS, the cell viability was calculated by a trypan blue assay (Gibco), and then placed on ice for subsequent use.

### scRNA-seq

Liver single-cell suspensions were loaded onto the 10x Genomics Chromium chip (10x Genomics; Pleasanton, CA, USA) to generate droplets. The obtained Gel Beads-in-emulsion was subjected to reverse transcription using a ProFlex PCR System (Thermo Fisher, Waltham, MA, USA). The resulting cDNA was purified and amplified. According to the cDNA concentration quantified by Qubit (Thermo Fisher), libraries were constructed with a Chromium Single Cell 3’ Library & Gel Bead Kit v3 (10x Genomics) following the manufacturer’s instructions. All libraries were sequenced by Novogene (Beijing, China) on the Illumina Novaseq platform.

### Histological analysis

Part of the flat tissue was cut from the liver tissue sample in the preservation solution and placed immediately in a 10% formalin solution. Tissue sections were cut at 5 μm thickness after being embedded in paraffin, followed by deparaffinization in xylene and rehydration in 100%, 95%, 90%, 80%, 75% alcohol successively. The sections were incubated with 3% H_2_O_2_, and nonspecific binding blocking was performed with 5% bovine serum albumin for 1 h. The sections were stained with hematoxylin and eosin (H&E) for histological evaluation and performed terminal deoxynucleotidyl transferase-mediated dUTP nick-end labeling (TUNEL) staining for apoptosis evaluation. Images were acquired on an Olympus BX63 microscope (Olympus, Shinjuku, Japan) at 200× magnification.

### Immunofluorescence staining

For immunofluorescence staining, liver sections were firstly incubated with primary antibodies against CD16 (1:500, Abcam, ab246222), FOS (1:500, Abcam, ab208942), SLC4A10 (1:50, Abcam, ab122229), CD3 (1:100, Abcam, ab16669), CD66b (1:500, Abcam, ab197678), S100A12 (1:500, Abcam, ab272713), KLRF1 (1:100, Proteintech, 21510-1-AP), Granzyme B (GZMB, 1:100, Invitrogen, MA1-80734), or Granzyme K (GZMK, 1:250, Abcam, ab282703) at 4°C overnight. After washing in 1× PBS with Tween 20, the appropriate fluorophore-conjugated secondary antibodies include Goat anti-Rabbit IgG H&L-Alexa Fluor 488 (1:100, Abcam, ab150077), Goat anti-Rabbit IgG H&L-Alexa Fluor 555 (1:100, Abcam, ab150078), and Goat anti-Rabbit IgG H&L-Alexa Fluor 647 (1:250, Abcam, ab150079) were used for incubation for 1.5 hours at room temperature. The antifade mounting medium with DAPI (Origene, Cat#ZU9557) was used for slides mounting. Fluorescence images of mucosal-associated invariant T (MAIT) cells, S100A12^+^ neutrophils, and FOS^+^ monocytes were captured with an Olympus BX63 microscope. Fluorescence images of granzyme B (GZMB)^+^ granzyme K (GZMK)^+^ natural killer (NK) cells were scanned by an Olympus SLIDEVIEW VS200 system.

### Data processing

Raw sequence files were processed with CellRanger 3.0.2 based on the GRCh38 reference for read alignment to generate the raw count data, which were further processed with Seurat,^18^ wherein cells with more than 4,000 unique features or with a mitochondrial percentage greater than 25% were filtered out to exclude doublets and dead cells. Gene symbols were revised according to the National Center for Biotechnology Information gene data (https://www.ncbi.nlm.nih.gov/gene/) updated on April 28, 2020, wherein unmatched genes and duplicated genes were removed.

### Cell type annotation

The raw data were normalized via the LogNormalize function. Principal component analyses (PCAs) were performed, followed by t-distributed stochastic neighbor embedding and uniform manifold approximation and projection (UMAP) analysis for dimensional reduction and clustering analysis. To annotate the cell type for each pre-computed cluster, scDeepSort^19^ and scCATCH^20^ were applied to obtain the predicted cell type for each cluster. Combined with the highly expressed genes and markers, each cluster was finally assigned a specific cell label.

### Single-cell trajectory analysis

Neutrophils were pre-processed and analyzed with monocle3^21^ with default parameters to generate the trajectory and dissect cellular decisions. The significantly correlated genes that cells use to navigate these decisions over pseudotime were tracked and ranked.

### Cellular module analysis

For each patient, modules consisting of GZMB^+^ GZMK^+^ NK cells, MAIT cells, and S100A12^+^ neutrophils before and after LT were extracted, respectively. The module score was defined as the total percentage of each cell type in each patient before and after LT.

### Cell-cell communication analysis

Cell-cell communication analysis was performed with scCrossTalk based on the highly expressed ligands of sending cells and receptors of receiving cells. For each cell type, human ligand-receptor pairs recorded in CellTalkDB^22^ were applied to filter out the significantly highly expressed ligands and receptors with a percentage of expressed cells > 25% and *P* < 0.05 using the *Z* score for each gene. For the ligand *L* of the sending cell type *i* and the receptor *R* of the receiving cell type *j,* the interacting score *S_(Li-Rj)_* was defined as the product of the average expression of *L* and *R*. A permutation test was then performed by randomizing the cell labels to re-calculate the interacting score. By repeating this step 1,000 times, the distribution *S* for the *L-R* interacting score between the *i* and *j* cells was obtained for comparison with the real interacting score, wherein the *P* value was calculated as follows:

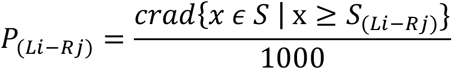

*L-R* pairs with a *P* value <0.05 were selected as the significantly enriched ligand-receptor interactions underlying the cell-cell communication between the pairwise sending cell type *i* and receiving cell type *j.*

### Pathway and biological process enrichment

The Metascape web tool (https://metascape.org/)^23^ was used to perform the enrichment of pathways and biological processes, wherein the top 100 highly expressed genes were selected according to the fold change of the average gene expression. Gene set enrichment analysis^24^ was performed using the ranked gene list with *clusterprofiler*^25^ to enrich the significantly activated pathways and biological processes, whose signatures were obtained from the Molecular Signatures Database v7.4 (MSigDB, http://www.gsea-msigdb.org/gsea/msigdb),^26^ including the gene sets from Gene Ontology (GO) and the canonical pathway gene sets derived from the Kyoto Encyclopedia of Genes and Genomes, Reactome, and WikiPathways pathway databases.

### Rat scRNA-seq data analysis

The processed scRNA-seq data matrix of six transplanted rat livers contained 23,675 cells involving 11 cell types.^15^ For NK cells, T cells, and granulocytes, the count data were normalized via LogNormalize, followed by PCA and UMAP analysis for dimensional reduction and clustering analysis. Pearson’s correlation coefficient was used to analyze the correlation of gene expression profiles between human and rat cell types, wherein the rat gene symbols were transformed to human gene symbols with the orthologs.

### Bulk RNA-seq data analysis

The raw bulk RNA-seq data matrix of liver cells from eight EAD and non-EAD patients after LT were downloaded from the Gene Expression Omnibus GSE23649 dataset, wherein gene expression values less than 0 were transformed to 0. The raw matrix was normalized by setting the median value to 1000 for each sample and transformed into a log2 matrix. To deconvolute the cell type composition of the bulk RNA-seq data, we used our scRNA-seq data containing 29 subtypes of 58,243 cells as the reference for RCTD,^27^ an R package for assigning cell types to bulk transcriptomics data; 100 representative cells for each subtype were randomly selected and applied in RCTD with all parameters kept as default.

### Statistical analysis

R (version 3.6.3) and GraphPad Prism 8.0.1 were used for the statistical analyses. Differences between two groups were determined using Welch’s t-test; *P* < 0.05 was considered to indicate a significant difference.

## RESULTS

### Overview of the single-cell transcriptomic atlas for human transplanted livers

A total of eight liver samples were collected from four donors and four recipients (Table 1) for scRNA-seq using 10x Genomics platform, and blood samples of recipients were collected for assessment of biochemical indicators (Supplementary Table S1), including alanine aminotransferase (ALT), aspartate aminotransferase (AST), total-bilirubin (T-Bil), and international normalized ratio (INR) (Fig. 1A). Histopathological evaluation with H&E and terminal deoxynucleotidyl transferase-mediated dUTP nick-end labeling (TUNEL) staining showed that liver injuries after LT were further aggravated compared with those before LT for these patients (Fig. 1B and Supplementary Fig. S1). According to the international definition of EAD,^6^ patient 2 was diagnosed as having EAD since the INR on postoperative day 7 was elevated at 1.74, whereas the other three patients were classified in the non-EAD group (Table 2).

**Fig. 1.**
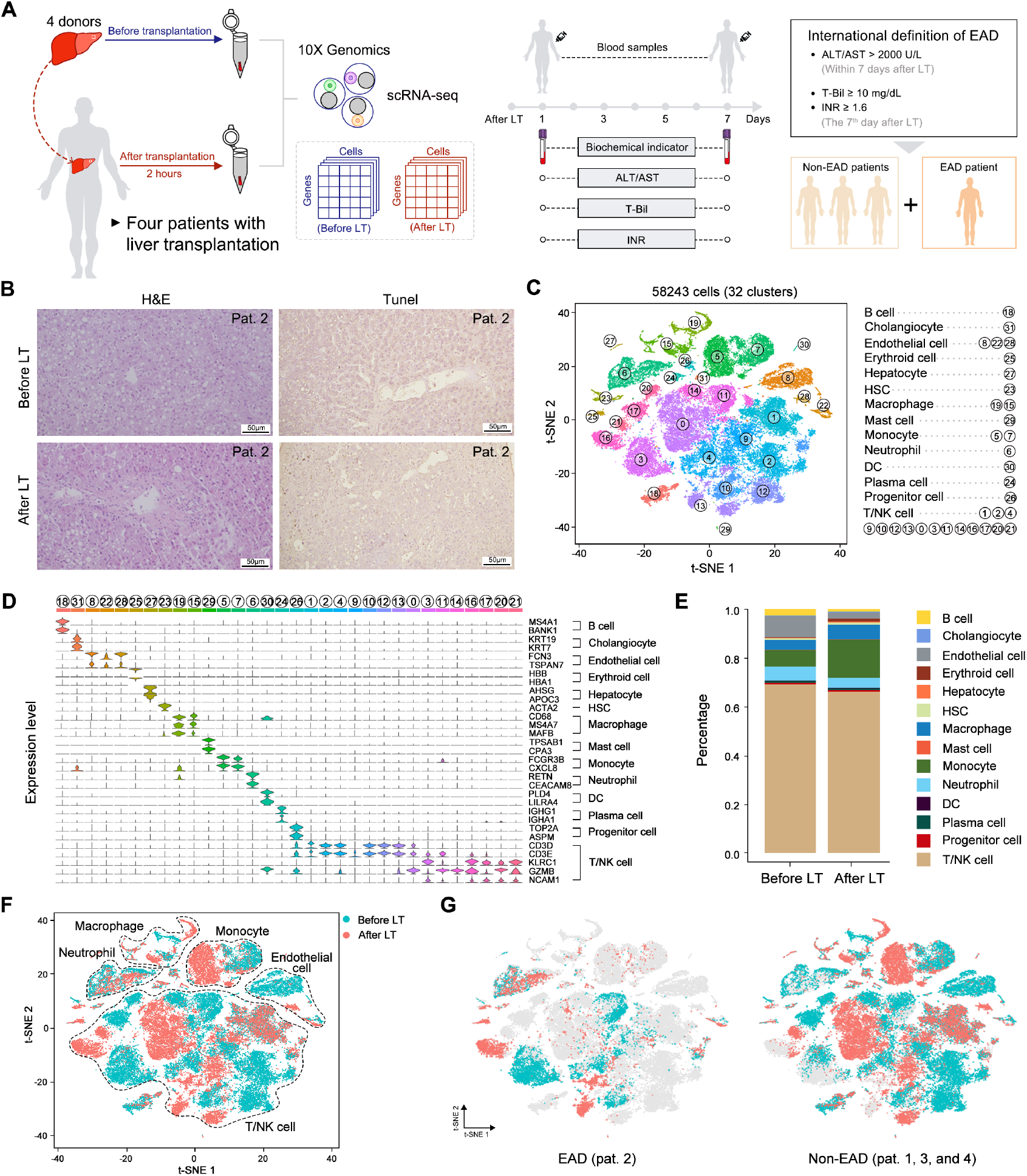
Single-cell transcriptomics atlas of human transplanted livers before and after LT. **(A)** An overview of the experimental design. Liver samples with cold perfusion and 2 hours after portal reperfusion were collected for scRNA-seq using the 10x Genomics platform followed by the collection of biomedical indicators with the first week. (**B)** H&E and Tunel staining of transplanted livers in patient 2 before and after LT. The scale bar = 50 μM. (**C)** The t-SNE plot of 32 cell clusters involving 58243 cells. (**D)** Violin plot of the representative marker genes for the 14 main cell types across 32 clusters. The expression level is the normalized count, namely the log1p value. (**E)** Ratio of cell number for the 14 main cell types among all liver cells before and after LT. (**F)** Distribution of endothelial cell, monocyte, macrophage, neutrophil, and T/NK cell before and after LT. (**G)** Difference of cells for the EAD and Non-EAD patients. The grey represents cells from other patients.

**Table 1.** Characteristics of donor livers and recipients.

| Characteristics | Donors (n = 4) |  |  |  |
| --- | --- | --- | --- | --- |
|  | 1 | 2 | 3 | 4 |
| Age | 40 | 20 | 64 | 32 |
| Sex | Male | Male | Female | Male |
| Graft weight (g) | 950 | 1000 | 1000 | 1800 |
| CIT (h) | 8 | ND | 7 | 8 |
| Viral hepatitis | No | No | No | No |
| Characteristics | Recipients (n = 4) |  |  |  |
|  | 1 | 2 | 3 | 4 |
| Age | 63 | 62 | 65 | 53 |
| Sex | Male | Male | Female | Male |
| BMI | 19.96 | 23.88 | 25.43 | 20.28 |
| High MELD | 39 | 36 | 40 | 39 |
| Operative time | 5h14min | 5h51min | 6h29min | 4h46min |
| Current state | Alive | Passed away | Alive | Alive |

CIT, Cold Ischemia Time; BMI, Body Mass Index; MELD, Model for End-stage Liver Disease; High MELD, preoperative MELD score >30.

**Table 2.**
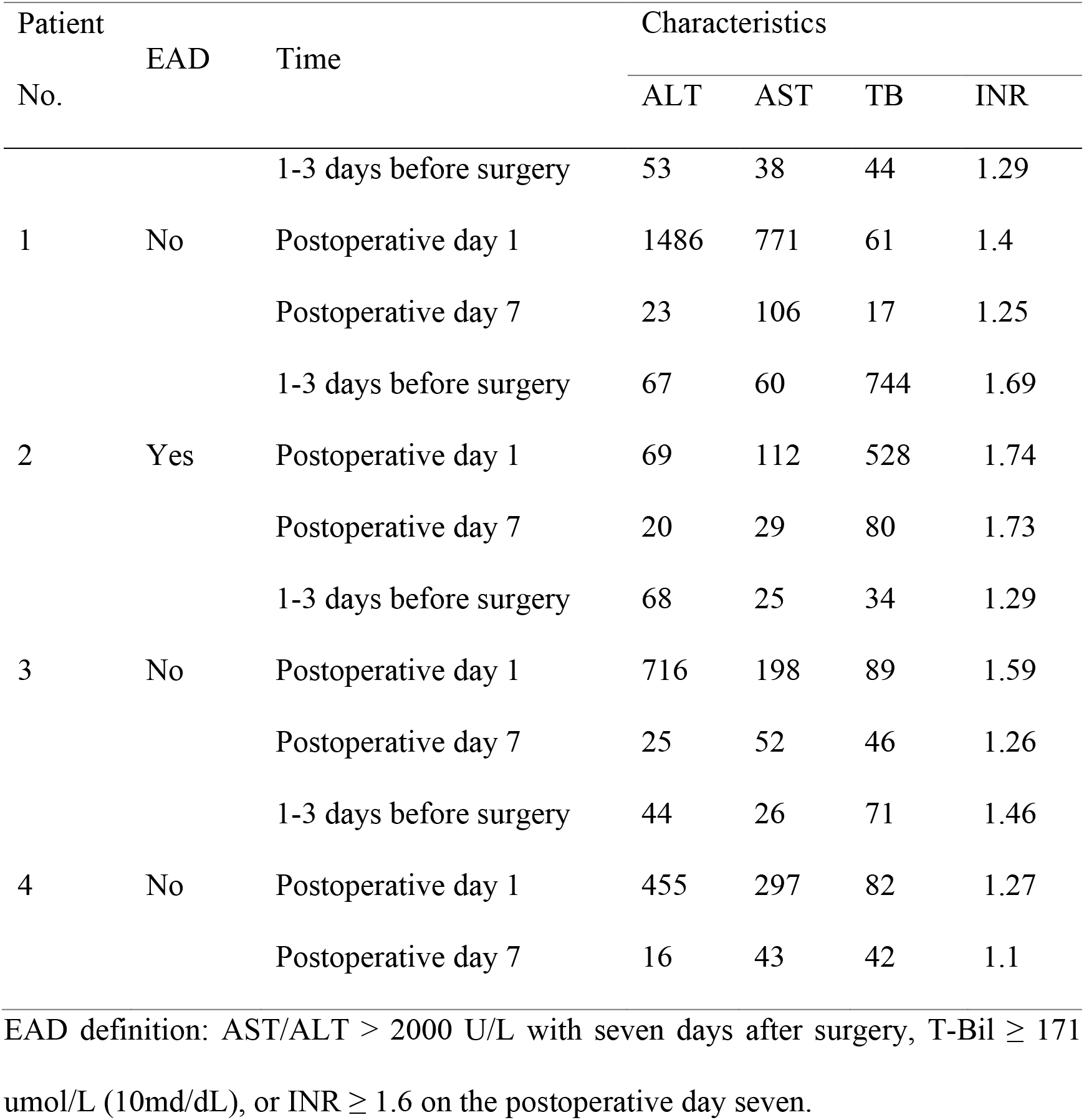
Biochemical parameters related to EAD definition before or after liver transplantation in four patients.

After quality control according to the number of unique features and mitochondrial percentage (Supplementary Fig. S2A), a total of 58,243 cells were included for further dimensionality reduction and clustering analysis (Supplementary Table S2), which generated 32 cellular clusters. Combining the predicted cell types of pre-trained scDeepSort^19^ and scCATCH^20^ with the highly expressed genes for each cluster, these 32 clusters were classified into 14 main cell types (Fig. 1C): B cells (MS4A1, BANK1), cholangiocyte (KRT7, KRT19), endothelial cells (FCN3, TSPAN7), erythroid cells (HBB, HBA1), hepatocytes (AHSG, APOC3), hepatic stellate cells (ACTA2), macrophages (CD68, MS4A7, MAFB), mast cells (TSPAB1, CPA3), monocytes (FCGR3B, CXCL8), neutrophils (RETN, CEACAM8), dendritic cells (PLD4, LILRA4), plasma cells (IGHG1, IGHA1), progenitor cells (TOP2A, ASPM), and T/ natural killer (NK) cells (CD3D, CD3E, KLRC1, GZMB, NCAM1), as shown in Figure 1D.

Compared to liver parenchymal cells, a substantial number of liver non-parenchymal cells were identified in the livers collected both before and after LT, which was attributed to the specific cell isolation protocol and kit used in our study. Specifically, T/ natural killer (NK) cells, neutrophils, macrophages, monocytes, and endothelial cells accounted for almost all of the cells obtained before and after LT (Fig. 1E). Unsurprisingly, these five main cell types exhibited either obviously different cell ratios or cell states after LT (Fig. 1F and Supplementary Fig. S2B), reflecting the fact that liver IR injuries are closely related to various types of inflammatory responses (e.g., immune cell activation, migration, and infiltration).^28, 29^ Notably, the EAD patient (patient 2) had a portion of T/NK cells with different cell states and a larger percentage of neutrophils after LT compared with those of the other three non-EAD patients (Fig. 1G), suggesting the crucial role of T/NK cell and neutrophil subtypes or substates during the graft remodeling after LT.

### High-resolution analysis of T/NK cell reveal unique cellular composition in patients after LT

According to the unique transcriptomic signatures, T/NK cells were divided into nine subtypes including mucosal-associated invariant T (MAIT), CD8^+^ GZMB^+^ T, CD8^+^ GZMK^+^ T, NKT, GZMB^+^ NK, GZMK^+^ NK, and GZMB^+^ GZMK^+^ NK cells (Fig. 2A). T cells, NK cells, and NKT cells are known to be involved in immune activation trigged by IR injury.^30^ Concordantly, CD8^+^GZMB^+^ T, GZMB^+^GZMK^+^ NK, and NKT cells all exhibited markedly increased proportions after LT (Fig. 2B). Notably, the proportion of MAIT cells decreased in the non-EAD patients (patients 1, 3, and 4) but increased in the EAD patient (patient 2) after LT (Fig. 2C, D and Supplementary Fig. S3). The proportion of GZMB^+^GZMK^+^ NK cells also remarkably increased in the EAD patient compared with that detected in the other, non-EAD, patients (Fig. 2C, E and Supplementary Fig. S4). Although a unique composition of T/NK cell subtypes was observed in patient 3 (non-EAD), largely contributing to the observed increases of CD8^+^GZMB^+^ T cells and NKT cells after LT (Fig. 2C), the gene expression profiles of these cells after LT resembled those of the T/NK subtypes before LT (Fig. 2F). However, GZMB^+^GZMK^+^ NK and MAIT cells after LT showed a dissimilar transcriptomic profile compared with those of the other subtypes before LT (Fig. 2F), suggesting specific functions of GZMB^+^GZMK^+^ NK cells and MAIT cells in transplanted livers.

**Fig. 2.**
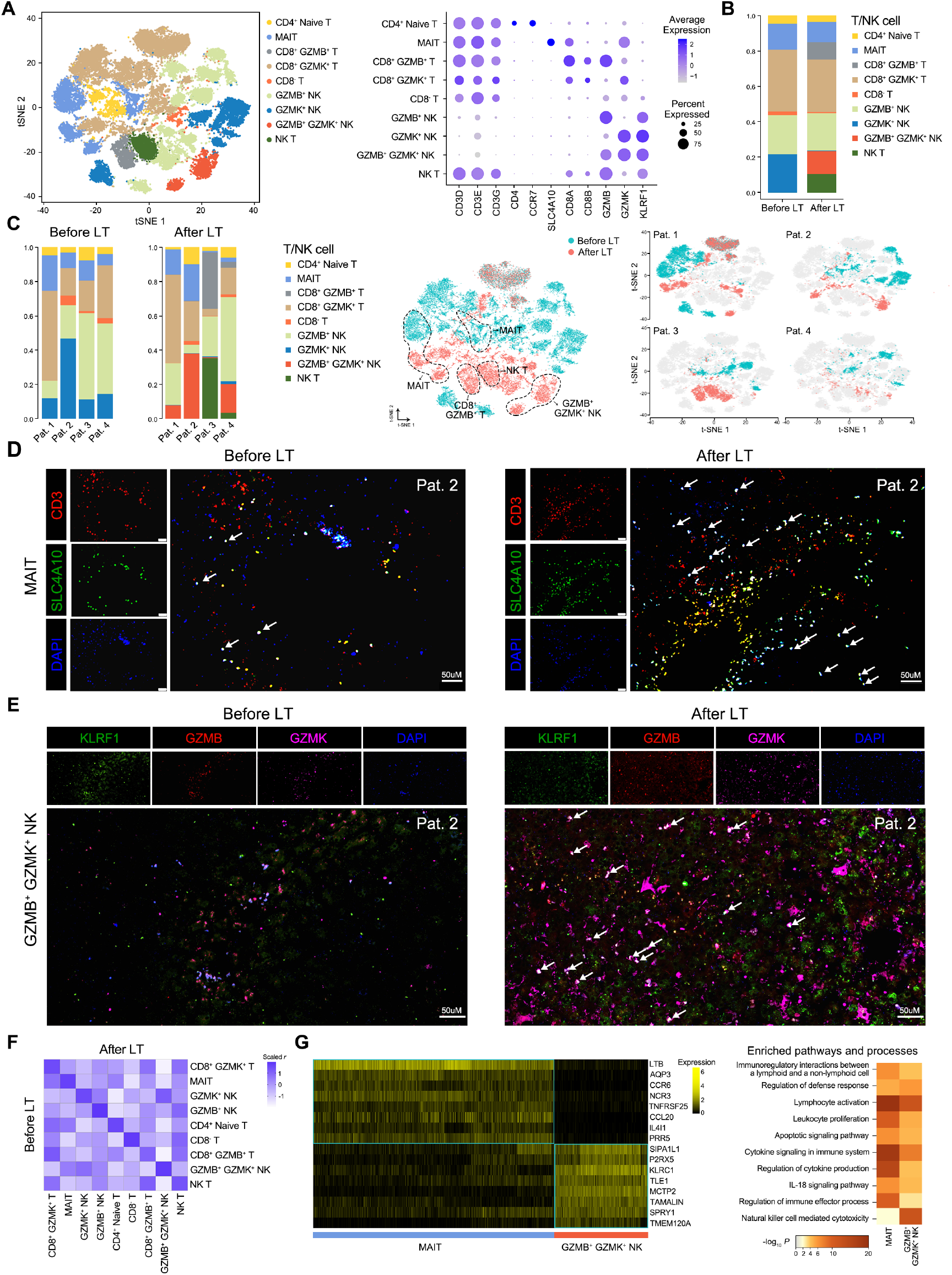
Distinct composition of T/NK cell subtypes in patients after LT. **(A)** Nine subtypes of T/NK cells (left). Known classical marker genes used to classify the nine subtypes (middle). (**B)** Ratio of cell number for the T/NK cell subtypes among T/NK cells before and after LT. (**C)** Ratio of cell number for the T/NK cell subtypes among T/NK cells across patients before and after LT. In the two-dimensional t-SNE plot, for each patient (right), the grey represents cells from other patients. (**D** and **E)** Immunofluorescent staining of MAIT cells (**D**) and GZMB^+^ GZMK^+^ NK cells (**E)** before and after LT. Scale bar = 50 μM. (**F)** Correlation of genome-wide expression profiling for nine T/NK subtypes before and after LT using the scaled Pearson’s coefficient. (**G)** Representative DEGs of GZMB^+^GZMK^+^ NK and MAIT cells and the top enriched pathways and processes according to the DEGs (*P* < 0.05). The expression level is the normalized count, namely the log1p value.

Next, we analyzed the highly expressed genes of GZMB^+^GZMK^+^ NK and MAIT cells as well as their enriched pathways and biological processes using Metascape^23^ to identify their specific functions (Supplementary Table S3). The genes upregulated in both of these cell types were significantly related with regulation of the defense response, lymphocyte activation, cytokine signaling in the immune system, and the IL-18 signaling pathway (Fig. 2G). Both MAIT and GZMB^+^GZMK^+^ NK cells were also prominently associated with immunoregulatory interactions between lymphoid and non-lymphoid cells (Fig. 2G), suggesting a vital role of these established injury-associated specific T/NK cell subtypes (MAIT and GZMB^+^GZMK^+^ NK cells) in the EAD patient via cell-cell interactions (CCIs) in the transplanted liver microenvironment.

### Key pro-inflammatory effect of S100A12^+^ neutrophils in patient 2 after LT

Given the enriched neutrophils in the EAD patient after LT (Fig. 1G), we further classified these cells into S100A12^+^ and S100A12^-^ neutrophils according to the expression level of S100A12, a protein highly associated with neutrophil activities^31^ (Fig. 3A). The increased S100A12^+^ neutrophils after LT were mainly distributed in patient 2, implying a specific distribution of postoperative S100A12^+^ neutrophils associated with EAD (Fig. 3B, C). To further confirm the state transition of S100A12^+^ neutrophils from S100A12^-^ neutrophils, single-cell trajectory analysis was performed with monocle3^21^ to dissect cellular decisions, demonstrating a single branch from the root (S100A12^-^ neutrophils) to the end state, namely S100A12^+^ neutrophils (Fig. 3D). Based on the reconstructed pseudotime trajectory, significantly correlated genes that cells use to navigate the decision over pseudotime were tracked and ranked, generating the top three pseudotime-associated genes: *S100A12, LTF,* and *PRTN3* (Fig. 3E). These findings are consistent with the differentially expressed genes (DEGs) identified between S100A12^+^ and S100A12^-^ neutrophils, wherein *S100A12* and *LTF* are markers of S100A12^+^ neutrophils and *PRTN3* is a marker of S100A12^-^ neutrophils (Fig. 3F and Supplementary Table S3).

**Fig. 3.**
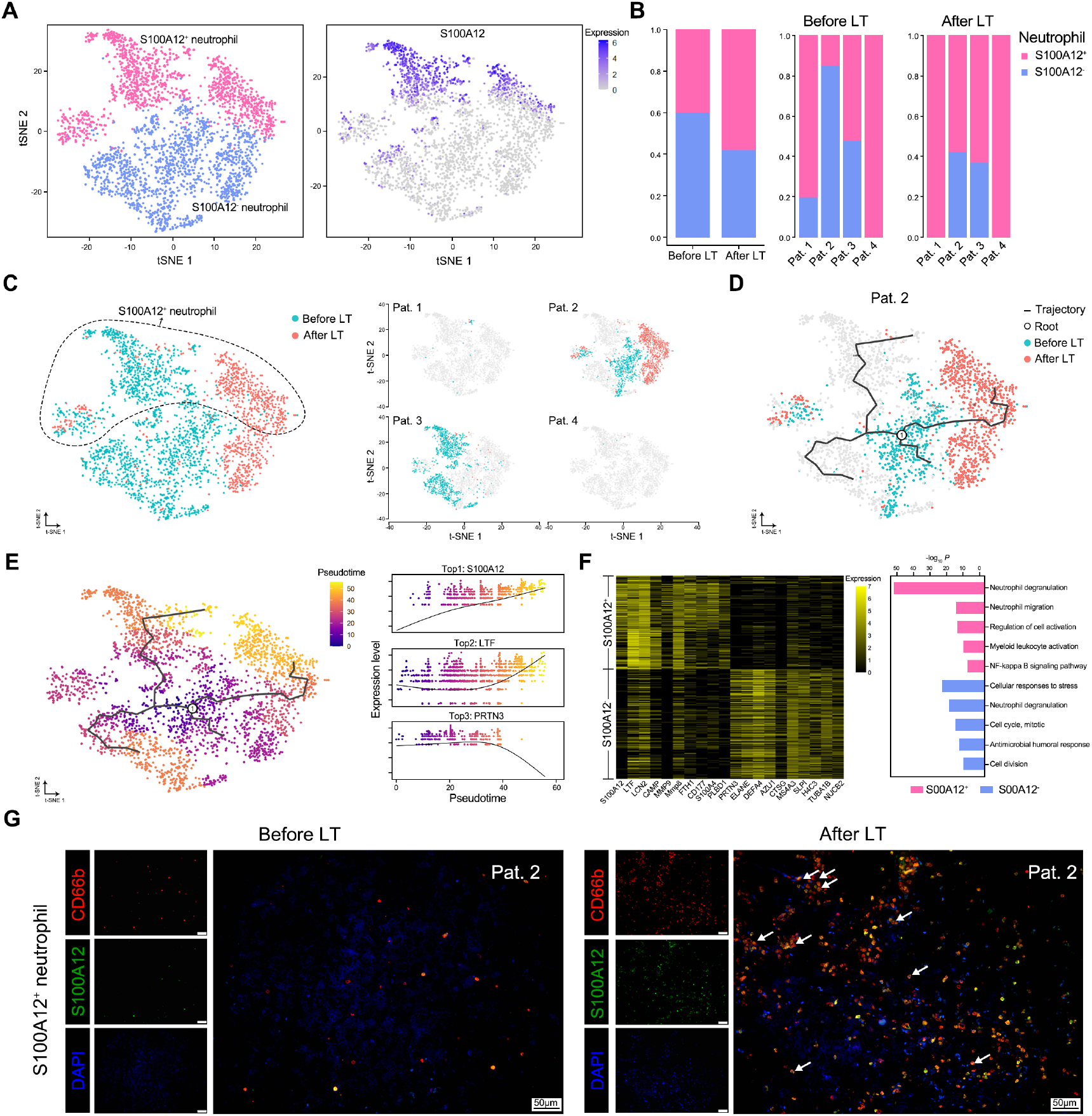
Function and trajectory analysis of neutrophils subtypes. (**A)** S100A12^+^ and S100A12^-^ neutrophil subtypes based on the expression of S100A12. The expression level is the normalized count, namely the log1p value. (**B)** Ratio of cell number for the two neutrophil subtypes among neutrophils. (**C)** Composition of two subtypes of neutrophils across four patients before and after LT. (**D)** Single-cell trajectory analysis of neutrophils before and after LT highlighted for pat. 2. (**E)** Pseudotime trajectory reconstruction and the top 3 significantly correlated genes that cells use to navigate the decision over pseudotime. (**F)** Representative DEGs of S100A12^+^ and S100A12^-^ neutrophil and the top enriched pathways and processes according to the DEGs (*P* < 0.05). (**G)** Immunofluorescent staining of S100A12^+^ neutrophils before and after LT. Scale bar = 50 μM.

Enrichment analyses showed that activated S100A12^+^ neutrophils are strongly related to neutrophil degranulation, migration, and the NF-kappa B signaling pathway, demonstrating distinct functions from S100A12^-^ neutrophils (Fig. 3F), consistent with the previous findings that S100A12, a damage-associated molecular pattern protein, is a sensitive marker for inflammation in various inflammatory disorders.^31^ Collectively, these results indicated that a non-negligible inflammatory response induced by increased S100A12^+^ neutrophils occurred in the transplanted liver of patient 2 (Fig. 3G and Supplementary Fig. S5), which may accelerate EAD progression.

### Subtype classification of mononuclear phagocytes and endothelial cells

Mononuclear phagocytes, comprising monocytes and macrophages, are major cell types in the immune remodeling of transplanted livers.^32^ Monocytes were divided into CD14^+^, CD16^+^, FOS^+^, and FOS^-^ monocytes (Supplementary Fig. S6A and Table S3). FOS^+^ monocytes showed a dramatic increase after LT (Supplementary Fig. S6B), which was most strongly detected in patient 3 (Supplementary Fig. S6C, D). FOS, also known as C-FOS, is a subunit of the AP-1 transcription factor complex, thereby promoting the transcription of genes encoding inflammatory mediators.^33^ The expression level of FOS is low in human resting monocytes and is increased in response to acute inflammatory stimulation,^33, 34^ indicating that activated monocytes-induced acute inflammation occurred in the transplanted liver of patient 3, whereas it seems to have no relationship with EAD. Macrophages were assigned to three subtypes, monocyte-derived macrophages, inflammatory Kupffer cells, and non-inflammatory Kupffer cells, with no obvious differences observed between the EAD patient and non-EAD patients (Supplementary Fig. S7A, B).

Endothelial cells were further classified into liver sinusoidal endothelial cells (LSECs) and vascular endothelial cells according to the transcriptome characteristics (Supplementary Fig. S8A and Table S3). The numbers of LSECs deceased after LT due to IR injuries. For each patient, LESCs occupied a small proportion both before and after LT, except for patient 4 (Supplementary Fig. S8B). These results indicated that graft remodeling is accompanied by damage of LSECs, accordant with the previous findings.^35, 36^

### Identification of a unique pathogenic cellular module associated with EAD

Compared with those of the non-EAD patients, the cellular module consisting of MAIT cells, GZMB^+^GZMK^+^ NK cells, and S100A12^+^ neutrophils demonstrated a specific and markedly increased tendency in the EAD patient (patient 2) after LT (Fig. 4A), and the highest module score was obtained for the EAD patient after LT (Fig. 4B). Therefore, we hypothesized that these three cell subtypes constitute a unique pathogenic cellular module related to the occurrence of EAD. Among the biochemical parameters, the EAD-associated pathogenic cellular module most highly corresponded to the level of INR (Fig. 4C). Furthermore, comparison of the DEGs in this pathogenic cellular module in the EAD patient and non-EAD patients showed that the EAD patient’s module after LT was significantly enriched in processes of neutrophil degranulation and migration, leukocyte cell-cell adhesion, and lymphocyte activation, which were not associated with the modules of non-EAD patients after LT (Fig. 4D and Supplementary Table S3).

**Fig. 4.**
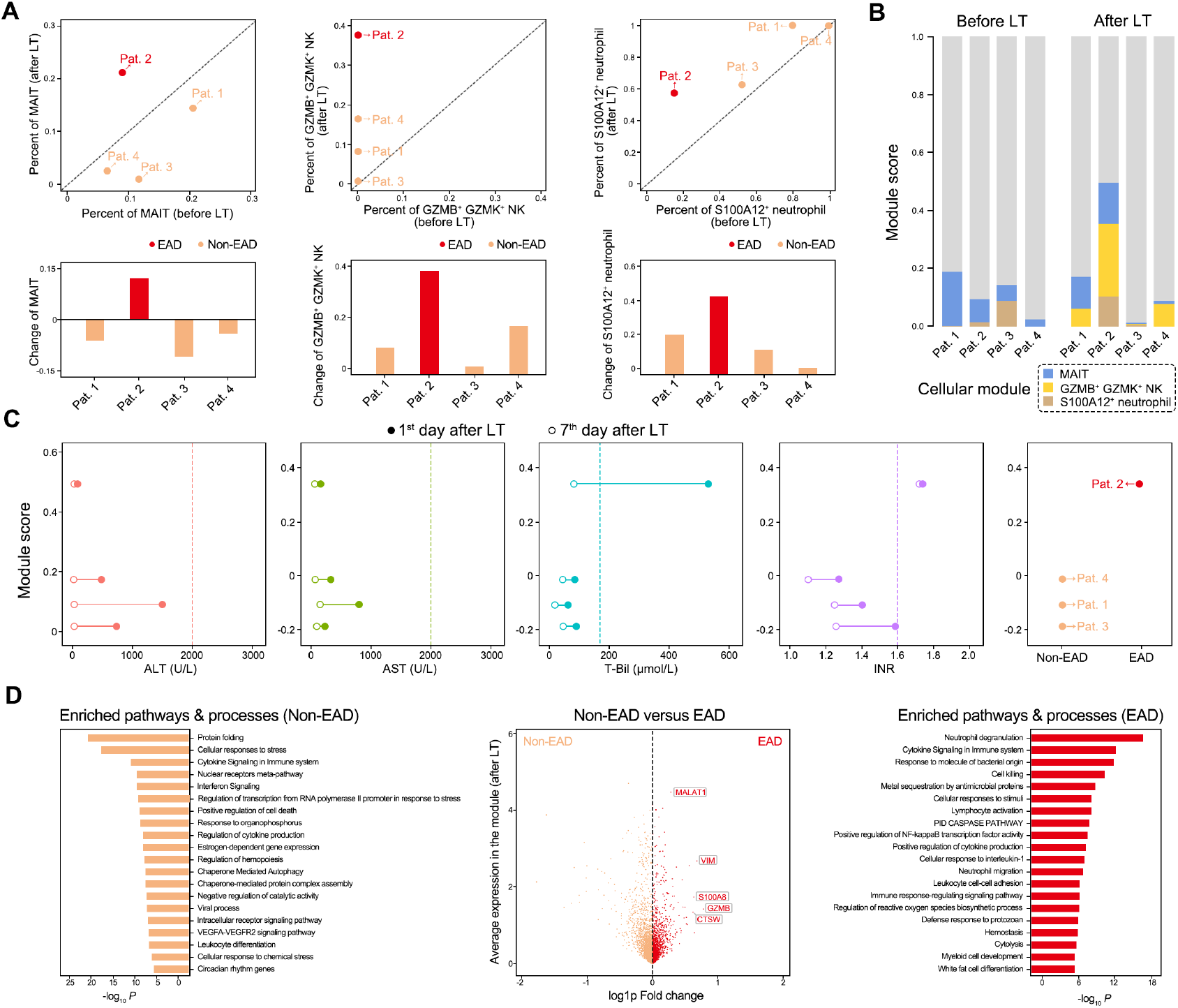
Association of the cellular module consisting of MAIT, GZMB^+^ GZMK^+^ NK cell, and S100A12^+^ neutrophil with EAD. (**A)** Percentage change of MAIT, GZMB^+^ GZMK^+^ NK cell, and S100A12^+^ neutrophil before and after LT. (**B)** Module score across four patients before and after LT. (**C)** Relationship of the cellular module with the biochemical indicators including the ALT, AST, T-Bil, and INR of four patients on the 1st and 7th day after LT. (**D)** DEGs of the cellular module in the EAD patient and the Non-EAD patients and the top enriched pathways and processes according to the DEGs (*P* < 0.05).

Given the crucial role of hepatocytes and LSECs in the maintenance of hepatic normal physiological functions and the finding of enriched immunoregulatory interactions between lymphoid and non-lymphoid cell, we next analyzed CCIs based on CellTalkDB^22^ between the pathogenic cellular module and hepatocytes as well as LSECs, integrated for the EAD patient and non-EAD patients, respectively (Fig. 5A). More ligand-receptor interactions were detected between GZMB^+^GZMK^+^ NK cells, MAIT cells, and S100A12^+^ neutrophils and hepatocytes/LSECs in the EAD patient than that in the non-EAD patients (Fig. 5B).

**Fig. 5.**
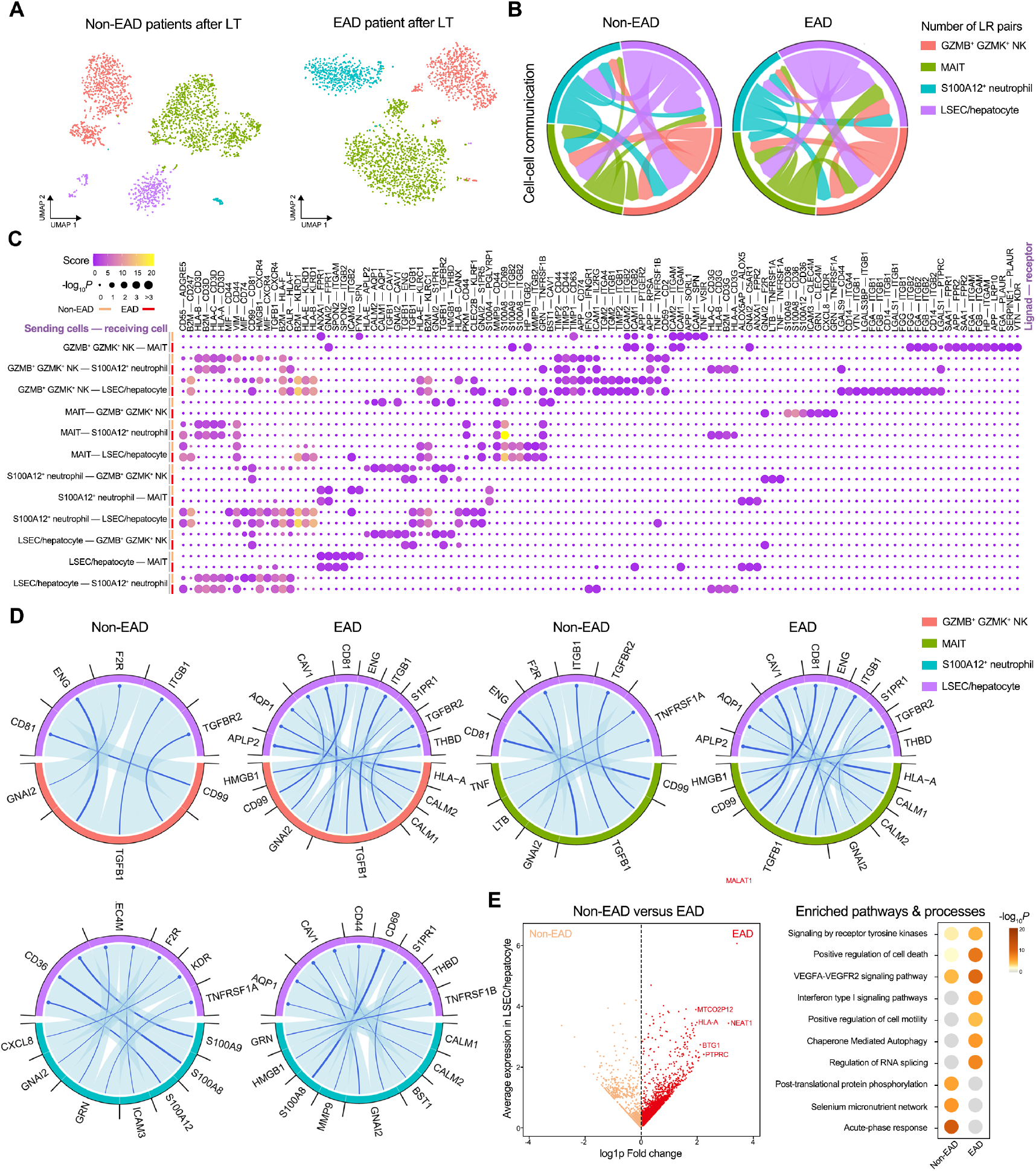
Cell-cell communications analysis of the cellular module. (**A)** Composition of the cellular module consisting of MAIT, GZMB^+^ GZMK^+^ NK cell, and S100A12^+^ neutrophil as well as LSEC/hepatocytes in the EAD and Non-EAD patients. (**B)** Number of enriched ligand-receptor pairs between pairwise cell types of MAIT, GZMB^+^ GZMK^+^ NK cell, S100A12^+^ neutrophil, and LSEC/hepatocyte (right). (**C)** Enriched ligand-receptor pairs between pairwise cell types. (**D)** Significantly enriched ligand-receptor pairs from the sending cell types in the module to the receiving cell types. (**E)** DEGs of the LSECs in the EAD patient and the Non-EAD patients and the top enriched pathways and processes according to the DEGs (*P* < 0.05).

In particular, the EAD patient showed specific CCIs among these cells (Fig. 5C) through several unique ligand-receptor pairs, which were the same for the interactions of GZMB^+^GZMK^+^ NK cells and MAIT cells with LSECs/hepatocytes (Fig. 5D). Four identical ligand-receptor pairs were found for CCIs from all three cell types to LSECs/hepatocytes in the EAD patient, demonstrating similar enhancement effects of this cellular module (Fig. 5D): HMGB1-THBD, CALM1-AQP1, GNAI2-S1PR1, and GNAI2-CAV1.

Finally, distinct differences in functions of LSECs/hepatocytes in EAD were identified according to the significantly upregulated DEGs between EAD and non-EAD patients (Supplementary Table S3), which were related to cell death, cell motility, and autophagy (Fig. 5E), indicating that the pathogenic cellular module may promote EAD by altering the states of LSECs/hepatocytes.

### Validation of the EAD-associated pathogenic cellular module

Due to the limited number of EAD cases, we further verified the pathogenic cellular module in two independent datasets. In the scRNA-seq dataset of six rat livers after LT^15^ (Fig. 6A), a cluster denoting *Gzmb^+^ Gzmk^+^* NK cells (cluster 2 of NK cell) was identified, and the gene expression profile strongly correlated with that of the GZMB^+^GZMK^+^ NK cells in our data (Fig. 6B and Supplementary Fig. S9A). The granulocyte subtype (cluster 2) specifically expressed the ortholog marker genes of human S100A12^+^ neutrophils, including *Lcn2, Camp9, Mmp9, Mmp8, S100a8,* and *S100a9* (Figs. 6C, 3F, and 5D). Moreover, the gene expression profiles of the granulocyte cluster 2 showed the strongest correlation with those of human S100A12^+^ neutrophils among all clusters in the rat data (Fig. 6D and Supplementary Figure S9B). Nevertheless, both GZMB^+^GZMK^+^ NK cells and S100A12^+^ neutrophils were present in the six rat livers after LT (Fig. 6E), indicating a common cellular mechanism across species.

**Fig. 6.**
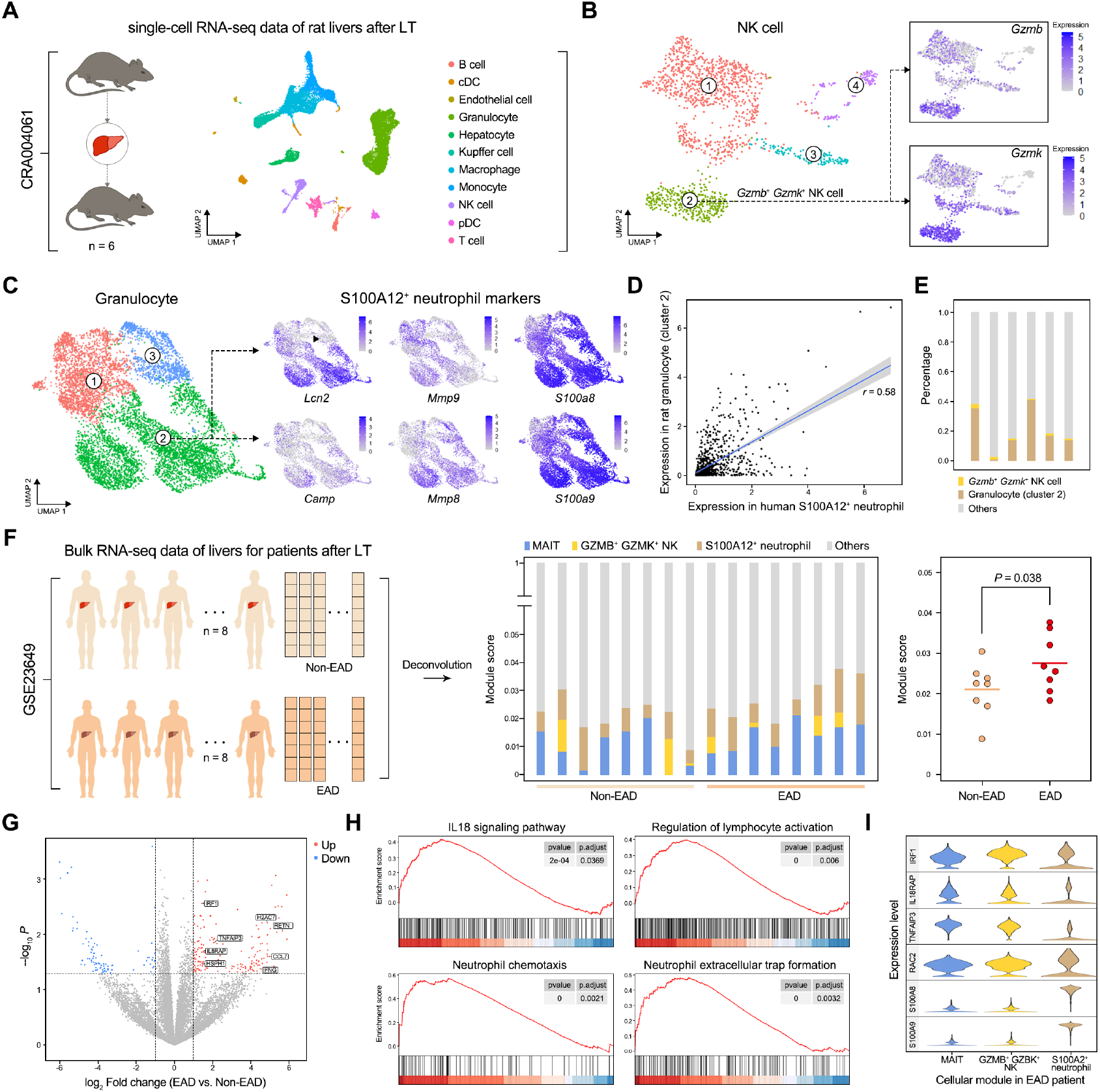
Validation of EAD-associated pathogenic cellular module. **(A)** Cell type composition of the single-cell RNA-seq dataset of six rat livers after LT. (**B)** Subtype analysis of *Gzmb^+^ Gzmk^+^* NK cells in rat liver. The expression level is the normalized count, namely the log1p value. (**C)** Subtype analysis of granulocytes in rat liver with known marker genes. (**D)** Correlation between human S100A12^+^ neutrophils and rat granulocytes cluster 2. (**E)** Ratio of cell number for GZMB^+^ GZMK^+^ NK cells and granulocytes cluster 2 among all cells for each rat. (**F)** Deconvolution of the bulk RNA-seq data of livers for 8 EAD and 8 Non-EAD patients after LT with RCTD by taking the scRNA-seq profiles of this study as the reference (left); The predicted percentage of the cellular module across 8 EAD patients and 8 Non-EAD patients (middle); Difference of the module percent between the EAD and Non-EAD patients after LT. *P* value was calculated with one-tailed Welch’s t test. (**G)** Volcano plot shows DEGs of the cellular module in the EAD and Non-EAD patients. The red represents up-regulated genes in EAD patients, while the blue represents up-regulated genes in Non-EAD patients. (**H)** Top enriched pathways and processes with Gene set enrichment analysis (GSEA, P < 0.05). (**I)** Expression level of genes related with top enriched pathways and processes across MAIT, GZMB^+^ GZMK^+^ NK cells, and S100A12^+^ neutrophils in our single-cell RNA-seq data.

Next, we analyzed another liver bulk RNA-seq dataset from eight EAD patients and eight non-EAD patients after LT (Fig. 6F). Deconvolution of bulk data with RCTD^27^ was used to obtain the proportions of each cell type. The pathogenic cellular module consisting of MAIT cells, GZMB^+^GZMK^+^ NK cells, and S100A12^+^ neutrophils was present in all patients after LT (Fig. 6F); however, the module scores in the eight EAD patients were significantly higher than those in the non-EAD patients, according to a one-tailed Welch’s t-test (Fig. 6F). Furthermore, the functions of upregulated genes in the eight EAD patients obtained by Gene Set Enrichment Analysis^24^ were related to the IL18 signaling pathway, lymphocyte activation, neutrophil chemotaxis, and neutrophil extracellular trap formation, in line with the enriched pathways of MAIT cells, GZMB^+^GZMK^+^ NK cells, and S100A12^+^ neutrophils from our single-cell data (compare Figs. 2G and 3F with Fig. 6G and 6H). Specifically, MAIT cells and GZMB^+^GZMK^+^ NK cells showed high expression of genes related to the IL18 signaling pathway and lymphocyte activation *(IRF1, IL18, RAP,* and *TNFAIP3),* whereas S100A12^+^ neutrophils showed high expression of *S100A8* and *S100A9* in our single-cell data (Fig. 6I), which are related to neutrophil chemotaxis and extracellular trap formation that were significantly activated in these eight EAD patients. These consistent results confirmed the high association of the unique pathogenic cellular module in transplanted livers after LT with EAD occurrence.

## DISCUSSION

In this study, we constructed the largest single-cell transcriptomic atlas of transplanted livers before and after LT, containing 58,243 liver cells from EAD and non-EAD patients and revealing a pathogenic cellular module consisting of MAIT cells, GZMB^+^GZMK^+^ NK cells, and S100A12^+^ neutrophils that is highly associated with the occurrence of EAD. This cellular module and its association with EAD occurrence were further verified in two independent LT datasets (rat and human).

The MAIT cells in the pathogenic module represent a novel and enriched population of innate immune cells in the human liver, which play complex roles in multiple liver diseases, including alcoholic/non-alcoholic/autoimmune liver disease, viral hepatitis, and liver cancer.^37, 38^ Specifically, MAIT cells play anti-bacterial roles in alcoholic liver disease, contribute to liver fibrosis by promoting hepatic stellate cell activation, and increase liver inflammation to further induce anti-inflammatory macrophage polarization. In this study, MAIT cells in transplanted livers were associated with immune regulation effects according to the high expression levels of associated genes such as *LTB, CCR6, IL4I1,* and *CCL20,* along with significant enrichment of lymphocyte activation, leukocyte proliferation, and cytokine production and signaling.

NK cells are the most abundant population of lymphocytes in the human liver, which are further recruited to the injury region to amplify the inflammatory response.^39, 40^ Although GZMB and GZMK are serine proteases known to be activated in NK cells, which could induce target cell apoptosis by caspase activation,^41, 42^ we identified a subtype of NK cells, GZMB^+^GZMK^+^ NK cells, in transplanted livers, which was more strongly associated with EAD occurrence compared to the GZMB^+^ NK or GZMK^+^ NK cells, suggesting the critical effects of GZMB^+^GZMK^+^ NK cells in liver graft remodeling.

S100A12^+^ neutrophils were also associated with the pathogenic EAD module. Human S100A12 is almost exclusively expressed and secreted by neutrophils and is dramatically overexpressed at inflammation sites.^43^ However, reports of S100A12 in liver disease mainly involve hepatocellular carcinoma and primary sclerosing cholangitis disease.^44, 45^ Herein, we identified S100A12^+^ neutrophils in transplanted livers, which may aggravate graft injury through pro-inflammatory effects.

The pathogenic cellular module identified in this study, including MAIT cells, GZMB^+^GZMK^+^ NK cells, and S100A12^+^ neutrophils, was also found in the two independent datasets, highlighting the universality of this module in transplanted livers across different species and platforms. The main difference was that MAIT cells were hardly observed in the rat transplanted livers (Supplementary Fig. S9C), which was not surprising as these cells are reported to be enriched in humans and much less abundant in murine species.^46, 47^ Moreover, the enriched pathways and biological processes of the pathogenic module were similar to that obtained from the independent human dataset with eight EAD patients, including lymphocyte and neutrophil activation as well as their associated immune response, inflammation, cytotoxicity, and tissue damage, which are reported as risk factors of EAD occurrence.^7, 31, 38, 41, 42^

Patient 2, who was the only patient diagnosed with EAD in our study based on the high level of INR, passed away two weeks after LT due to multiple organ failure. As a prothrombin time-related indicator, INR is used to evaluate the severity and prognosis of acute liver failure, and its increase is a critical feature of advanced liver failure as prothrombin time will not be extended until 80% of synthetic ability is lost.^48^ Collectively, our results suggest that the EAD-associated pathogenic cellular module might mainly contribute to prothrombin-associated liver injury in EAD, whereas it needs further exploration.

Overall, we have proposed a pathogenic cellular module associated with the occurrence of EAD after LT at the single-cell resolution, providing new insights into the understanding of EAD. Intervention in this pathogenic cellular module might be a novel direction for preventing the occurrence of EAD.

## DATA AND CODE AVAILABILITY

The accession number for the raw and processed counts data reported in this paper is Gene Expression Omnibus (GEO, https://www.ncbi.nlm.nih.gov/geo/): GSE189539. The bulk RNA-seq data of liver samples for eight EAD and non-EAD patients after LT were collected from GEO: GSE23649. The scRNA-seq data of six rat transplanted livers are accessible at Genome Sequence Archive (https://ngdc.cncb.ac.cn/gsa/): CRA004061. Analysis scripts for the scRNA-seq data processing pipeline are available at the Satija Lab tutorial (https://satijalab.org/seurat). Source codes for the scCrossTalk R package are provided on GitHub (https://github.com/ZJUFanLab/scCrossTalk).

## ACKNOWLEDGMENTS

This work was financially supported by the National Natural Science Foundation of China (81973701, 81903767), the Natural Science Foundation of Zhejiang Province (LZ20H290002), the Innovation Team and Talents Cultivation Program of National Administration of Traditional Chinese Medicine (No: ZYYCXTD-D-202002), the Key Program, National Natural Science Foundation of China (81930016), the National Key R&D Program of China (2021YFA1100500), and the China Postdoctoral Science Foundation (2021M702828). We thank the Novogene Biotechnology Co., LTD (Tianjin, China) and Morphological Platform of Zhejiang University School of Medicine (Hangzhou, China) for their technical support.

## AUTHOR CONTRIBUTIONS

Z.W. designed, performed, and analyzed all experiments; X.S. processed scRNA-seq data and performed computational analysis; K.W. and P.Z. participated in the experiment; Z.W. and X.S. wrote the manuscript; X.Y. provided the scRNA-seq matrix of rat livers. X.L., L.Z., S.Z., X.X., and X.F. supported and supervised the experiment and revised the manuscript; X.F. and X.X. conceptualized the study. All the authors reviewed the manuscript.

## COMPRTING INTERESTS

The authors declare no competing interests.

---

**Fig. S1.**
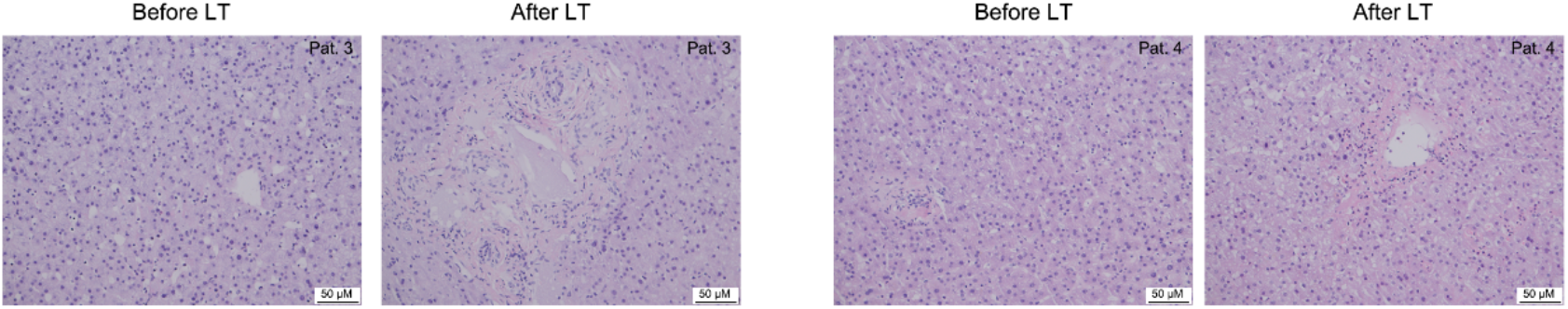
Histological staining of transplanted livers of other patients. H&E staining was performed before and after LT. The scale bar = 50 μM.

**Fig. S2.**
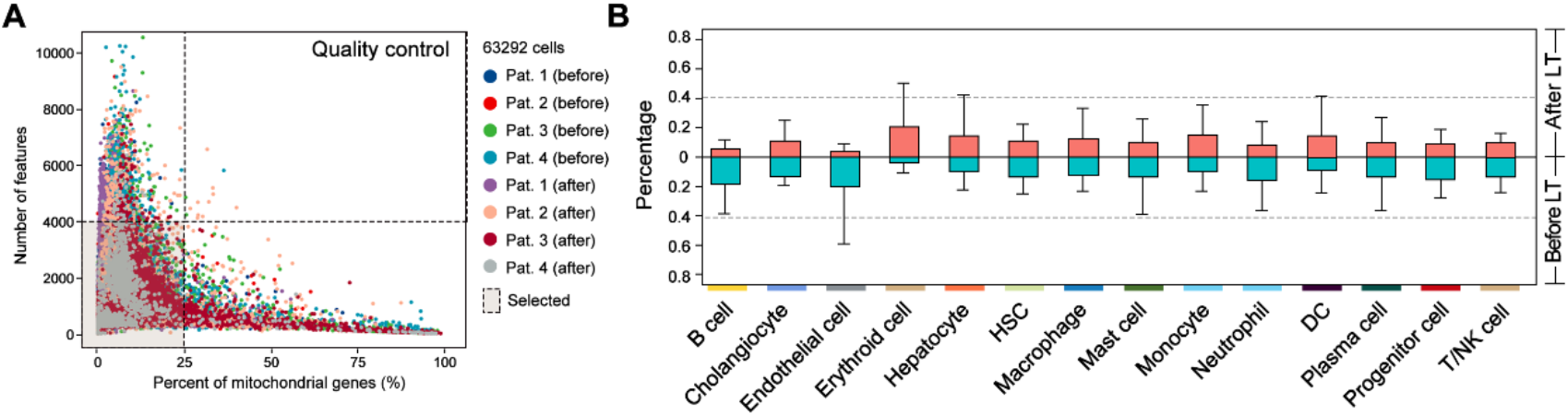
Single-cell transcriptomic atlas of human transplanted livers. **(A)** Quality control with the number of unique features (< 4000) and mitochondrial percent (< 25%). **(B)** The bar plot of the percentage of each cell type in each sample across four patients before and after LT, respectively, representing mean ± sd.

**Fig. S3.**
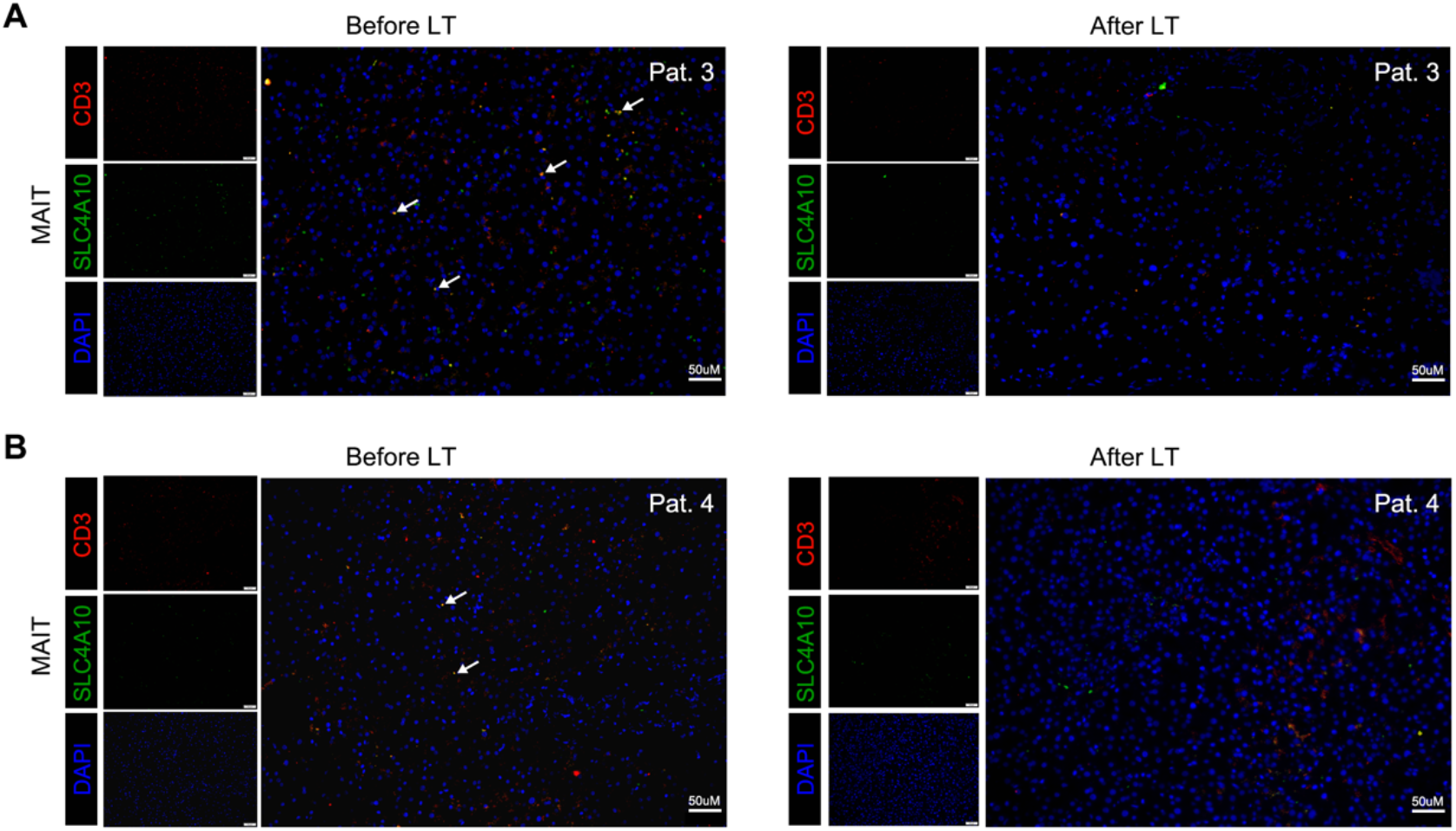
Validation of MAIT cell of other patients before and after LT. Immunofluorescent staining of MAIT cells before and after LT in patient 3 **(A)** and patient 4 **(B)**. Scale bar = 50 μM.

**Fig. S4.**
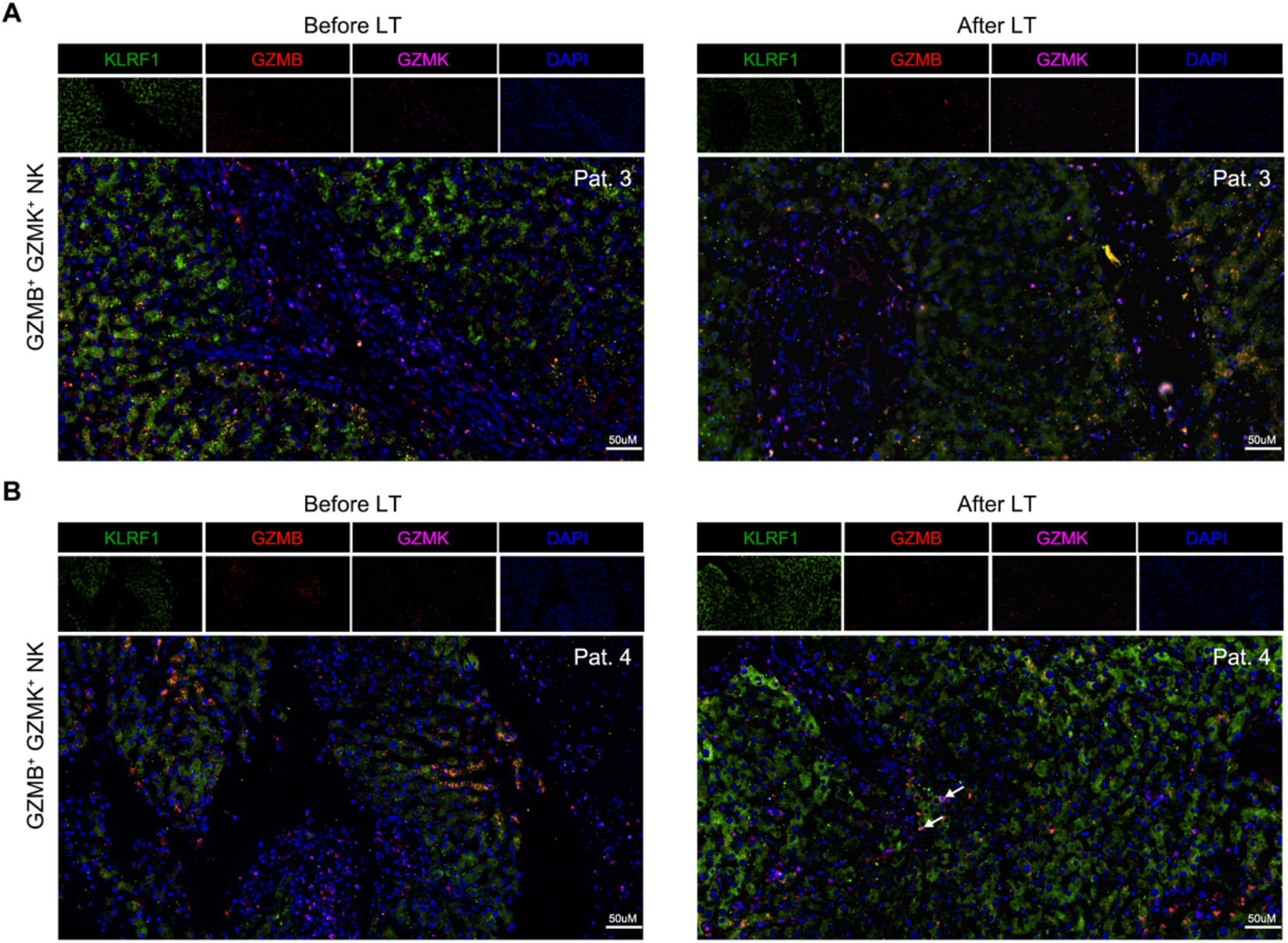
Validation of GZMB^+^GZMK^+^NK cell of other patients before and after LT. Immunofluorescent staining of GZMB^+^ GZMK^+^ NK cells before and after LT in patient 3 **(A)** and patient 4 **(B)**. Scale bar = 50 μM.

**Fig. S5.**
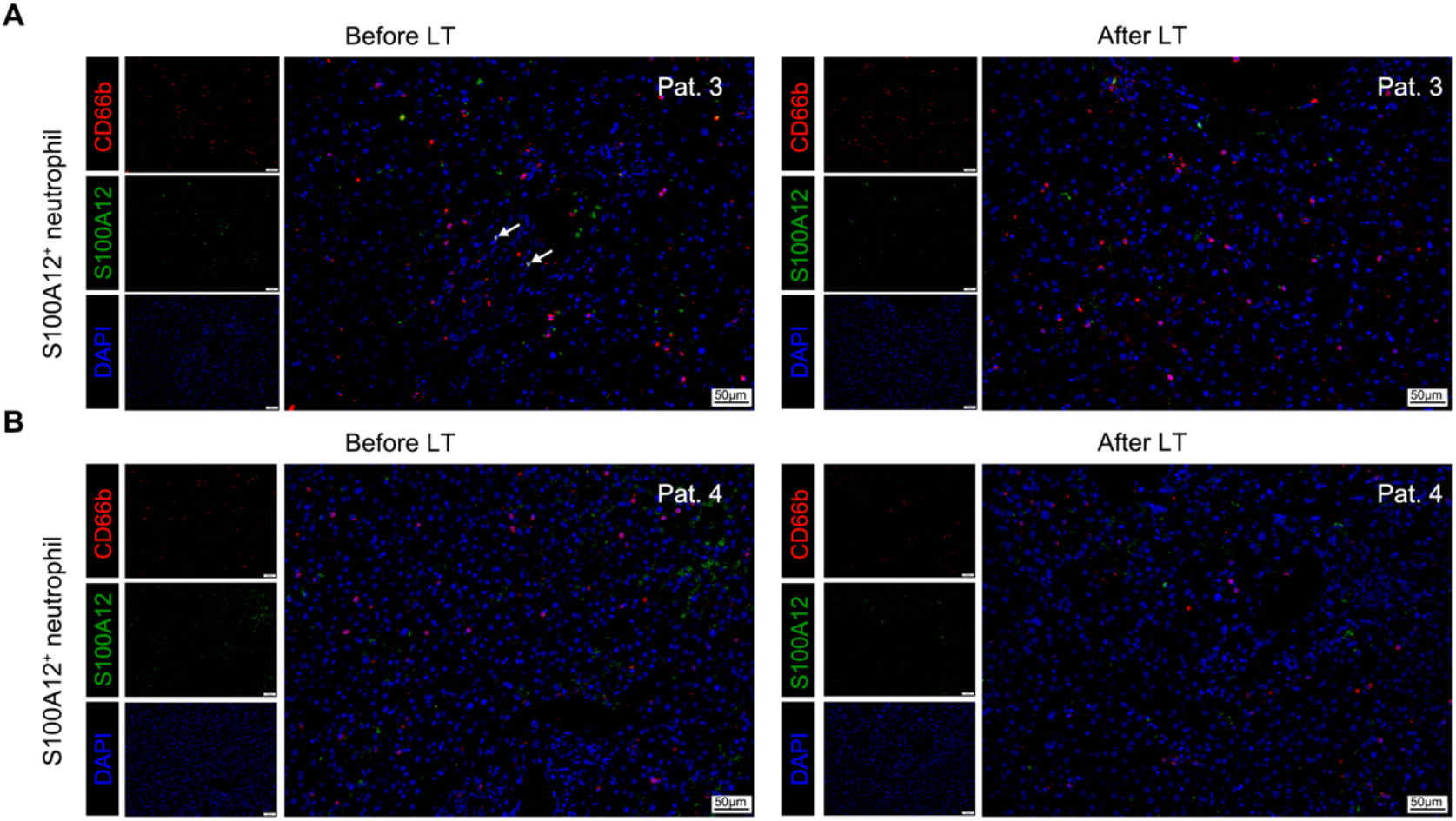
Validation of S100A12^+^ neutrophils of other patients before and after LT. Immunofluorescent staining of S100A12^+^ neutrophils before and after LT in patient 3 **(A)** and patient 4 **(B)**. Scale bar = 50 μM.

**Fig. S6.**
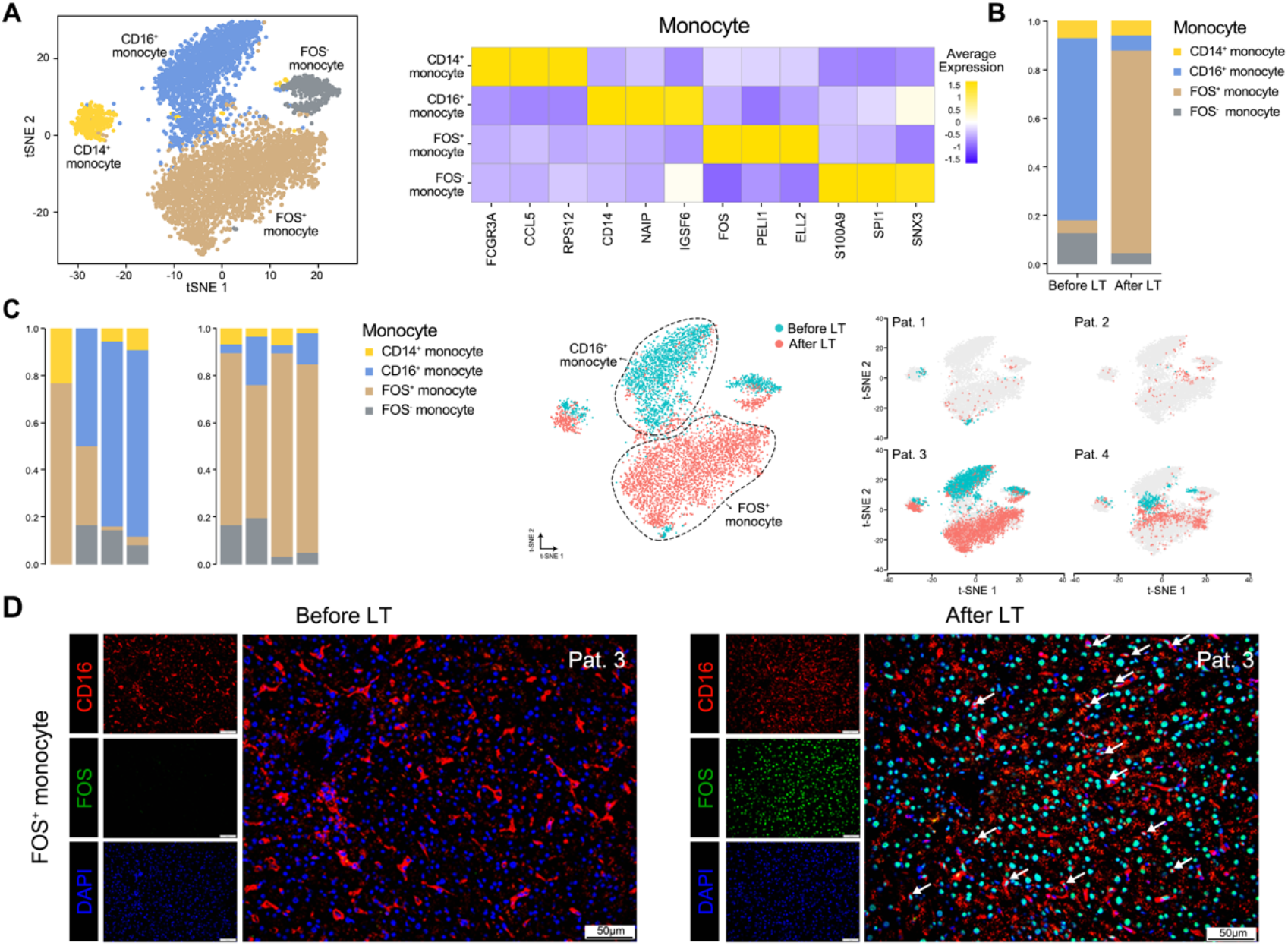
Analysis of cell subtypes of monocytes. **(A)** Four subtypes of monocytes, namely CD14+, CD16+, FOS+, and FOS-monocytes (left). Known marker genes, e.g., CD14, FCGR3A, and FOS, used to annotate the four subtypes. The color represents the scaled average gene expression level in each subtype (right). **(B)** Ratio of cell number for the four monocyte subtypes before and after LT. **(C)** The composition of four subtypes of monocytes across four patients before and after LT. In the two-dimensional t-SNE plot, the blue represents cells before LT, while the red represents cells after LT. For each patient (right), the grey represents cells from other patients. **(D)** Immunofluorescent staining of FOS+ monocytes before and after LT. Liver sections were stained with antibodies against CD16 and FOS; DAPI was used for nucleus staining; the scale bar = 50 μM.

**Fig. S7.**
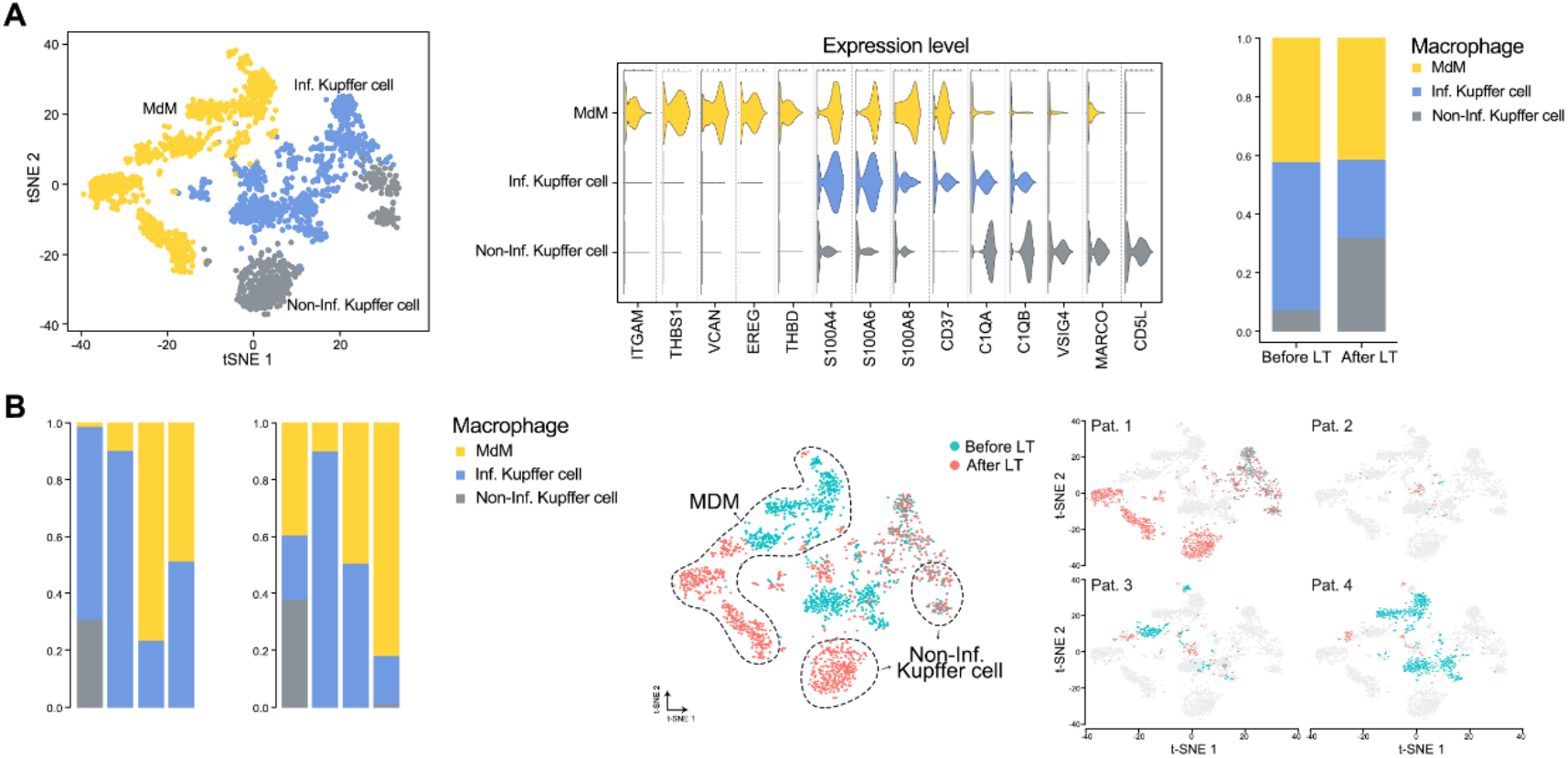
Analysis of cell subtypes of macrophages. **(A)** Three subtypes of macrophages, namely monocyte-derived macrophage (MdM), Inflammatory (Inf.) Kupffer cell, and Non-inflammatory (Non-Inf.) Kupffer cell. Known marker genes for three subtypes of macrophages (middle). The expression level is the normalized count, namely the log1p value. Ratio of cell number for the three macrophage subtypes before and after LT (right). **(B)** The composition of three subtypes of macrophages across four patients before and after LT. In the two-dimensional t-SNE plot, the blue represents cells before LT, while the red represents cells after LT. For each patient (right), the grey represents cells from other patients.

**Fig. S8.**
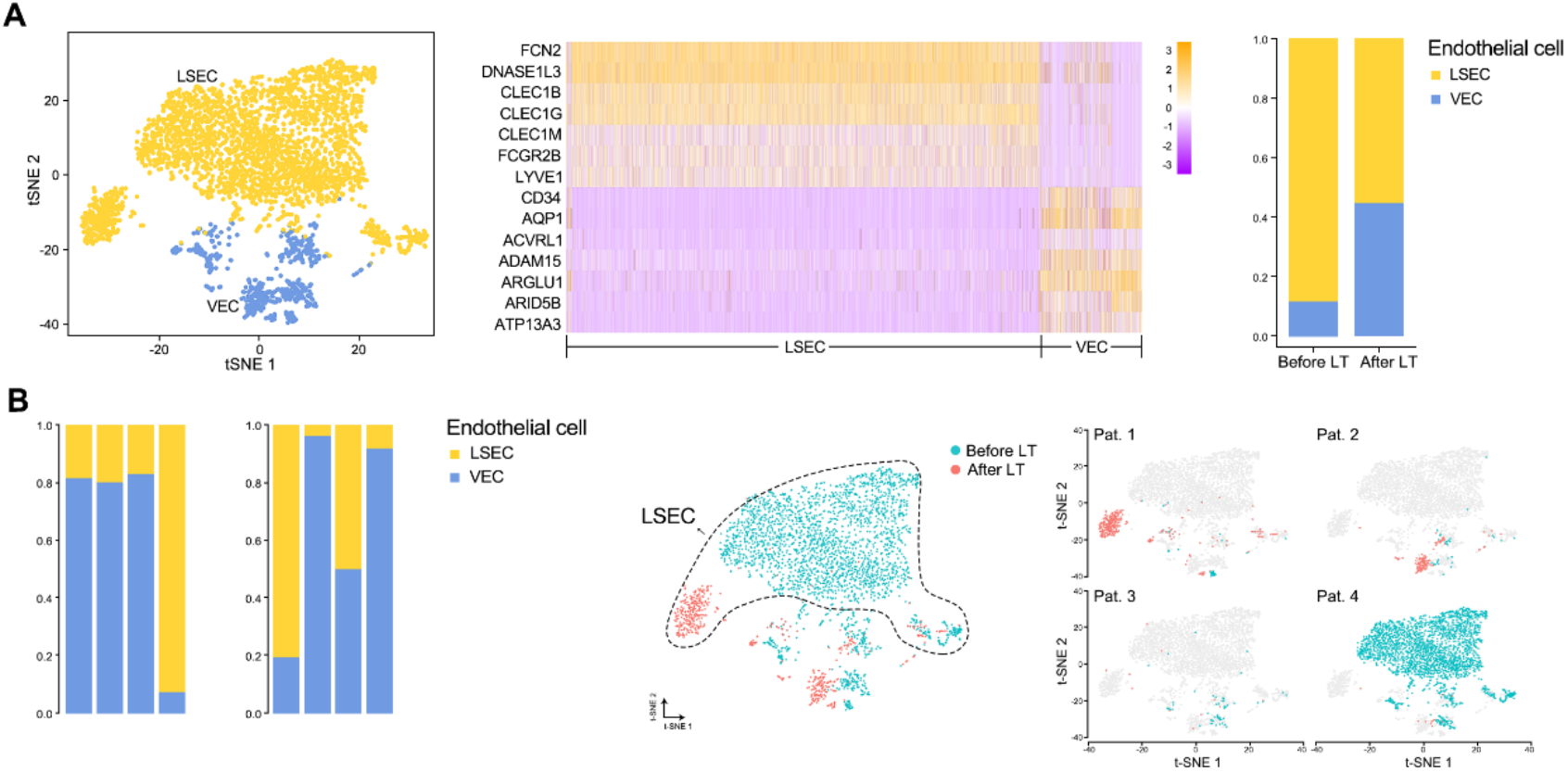
Analysis of cell subtypes of endothelial cells. **(A)** Two subtypes of endothelial cells, namely liver sinusoidal endothelial cells (LSECs) and vascular endothelial cells (VECs). Known marker genes for two subtypes of endothelial cells (middle). The expression level is the normalized count, namely the log1p value. Ratio of cell number for the two endothelial cell subtypes before and after LT (right). **(B)** The composition of two subtypes of endothelial cells across four patients before and after LT. In the twodimensional t-SNE plot, the blue represents cells before LT, while the red represents cells after LT. For each patient (right), the grey represents cells from other patients.

**Fig. S9.**
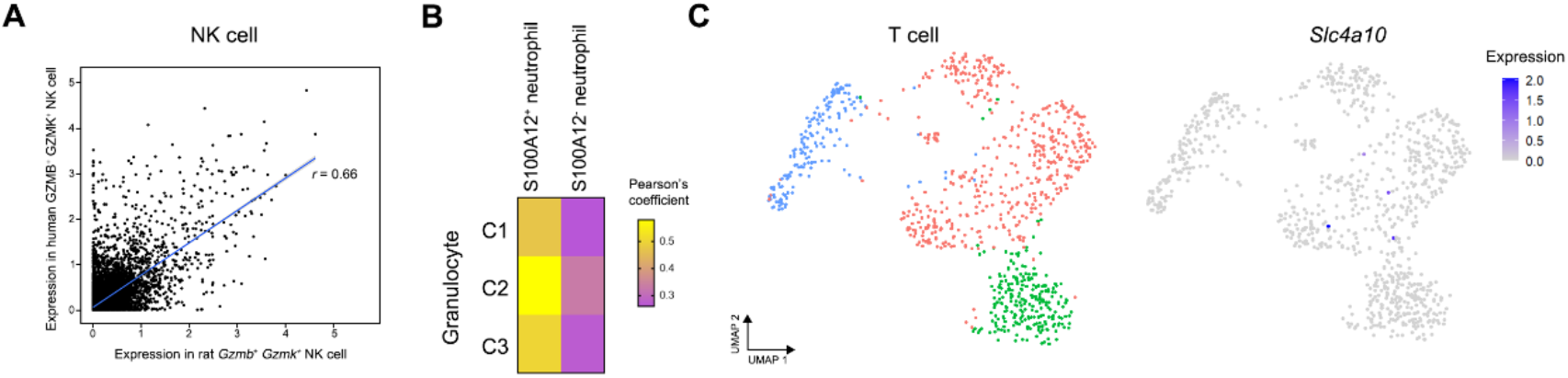
Validation of the EAD-associated pathogenic cellular module. **(A)** Correlation between human GZMB^+^ GZMK^+^ NK cells and rat *Gzmb^+^ Gzmk^+^* NK cells. **(B)** Clustering analysis of rat T cells and the gene expression of MAIT cell marker *Slc4a10.* The expression level is the normalized count, namely the log1p value. **(C)** Pearson’s correlation coefficients between human S100A12^+^/S100A12^-^ neutrophils and three clusters of rat granulocytes.

